# Senescent cells exhibit features of developmental signaling centres

**DOI:** 10.1101/2025.10.30.685565

**Authors:** Daniel Sampaio Gonçalves, Ludivine Dulac, Birgit Ritschka, Muriel Rhinn, Annabelle Klein, Christopher D.R. Wyatt, Manuel Irimia, William M. Keyes

**Affiliations:** Institut de Génétique et de Biologie Moléculaire et Cellulaire (IGBMC), Équipe Labellisée Ligue Contre le Cancer, Illkirch, France; UMR7104, Centre National de la Recherche Scientifique (CNRS), Illkirch, France; U1258, Institut National de la Santé et de la Recherche Médicale (INSERM), Illkirch, France; Université de Strasbourg, Illkirch, France; Centre for Genomic Regulation (CRG), The Barcelona Institute of Science and Technology, Dr. Aiguader 88, Barcelona 08003, Spain; Universitat Pompeu Fabra (UPF), Barcelona, Spain; ICREA, Pg. Lluis Companys 23, Barcelona 08010, Spain

**Keywords:** senescence, SASP, development, aging, signaling centre, organizer, muscle, cancer

## Abstract

Cellular senescence is a state that contributes to tissue patterning during development and tissue repair, but which becomes misregulated in ageing and disease. The senescence-associated secretory phenotype (SASP) mediates many of the context-dependent effects, yet its molecular composition and evolutionary origins remain incompletely understood. Here, we characterize the SASP signature of developmental senescent cells and find that it is enriched for genes encoding major developmental signaling pathways. Integrative analyses of *in vitro* and *in vivo* transcriptomes reveal that human and mouse senescent cells across diverse contexts consistently activate these developmental programs, with key morphogens constituting conserved SASP components. Functionally, we show that the SASP is sufficient to induce the expression of central developmental genes and transcription factors across multiple cell types. These findings show that senescent cells resemble developmental signaling centers and uncover developmental signaling reactivation as an evolutionarily conserved feature of senescence, providing conceptual insight into its physiological functions and its pathological impact during ageing.

## INTRODUCTION

Cellular senescence is a state of stable cell cycle arrest accompanied by a robust secretory activity, known as the senescence-associated secretory phenotype (SASP)^1,2^. Senescence is closely associated with aging and tumor suppression, functioning as a barrier to uncontrolled proliferation.^3,4^ Furthermore, the accumulation of senescent cells is implicated in various pathological conditions, including neurodegeneration, fibrosis, and cancer, where their persistent presence exacerbates tissue dysfunction and promotes chronic inflammation^5–7^. In these situations, senescent cells can exert multifaceted effects on neighboring cells, including promoting complex gene-expression and cellular changes.

However, senescence is not exclusively detrimental. The transient appearance of senescent cells plays beneficial roles following tissue injury and wounding, including in the lung, skin and heart ^8–10^. Senescent cells can also promote cell plasticity and reprogramming, and contribute to tissue regeneration^11–13^. In addition, they have demonstrated roles during embryonic development, contributing to tissue morphogenesis and fate-instruction^14,15^. These beneficial associations have suggested an evolutionary origin for senescence as initially a beneficial process, that became misregulated during aging and disease^5,16^.

During embryogenesis, features of senescence have been identified in some developmental signaling centers, such as the apical ectodermal ridge (AER) in the limb and in the hindbrain roof plate^14^. These structures display hallmark properties of senescence, including SA-β-gal activity, p21 expression, low proliferative index, and a high secretory profile. Indeed the latter is a hallmark feature of developmental signaling centres, or organizers, which by definition are cell populations that direct cell fate specification and tissue morphogenesis via paracrine signaling, contributing to tissue patterning and embryo development^17–19^. Signaling centers orchestrate development through the secretion of conserved morphogens such as BMP’s, FGF’s, WNT’s, IGFBP’s and other factors, often in spatial gradients that pattern adjacent tissues.

Recently, we performed RNA sequencing profiling of the p21-positive senescent AER, to identify a gene-expression signature of developmental senescence^20^. From this, we identified that there is a significant overlap in genes expressed in developmental senescence, and senescence in other contexts. Given these associations, this raises an interesting conceptual question: As senescent cells and some signaling centres share many features, including SA-β-gal, p21, and SASP, might senescent cells exhibit additional properties associated with developmental signaling centres?

Here, we characterize the SASP signature of developmental senescent cells and find that they express genes encoding key developmental signaling pathways, including BMP, FGF, IGFBP, and WNT families, among others. Comparison with RNA-sequencing datasets from in vitro and in vivo models reveal that senescent cells across diverse contexts consistently upregulate similar developmental pathways, with core morphogens emerging as common SASP components. Functionally, we show that the SASP from senescent cells can induce the expression of pivotal developmental genes and transcription factors in multiple cell types. Together, these findings uncover that senescent cells exhibit properties of developmental signaling centres and suggest that the reactivation of developmental signaling is a conserved feature of senescence, providing insight into its physiological roles and its pathological consequences when misregulated.

## RESULTS

### Developmental signaling factors are prominent features of developmental senescence

Recently we performed RNA sequencing to explore the gene expression signature of developmental senescent cells^20^. Using a newly-generated p21-mCherry-CreERT2 mouse model, we isolated the p21-positive AER, and compared its signature to the adjacent non-senescent ectoderm (Fig. 1a,b). As the entire AER is p21 and SA-β-Gal positive at this developmental stage, this analysis provides a signature of both the AER and developmental senescence – the two definitions refer to the same population of cells. This uncovered a gene-expression signature of developmental senescence that we demonstrated is conserved across many senescent states^20^.

**Figure 1.**
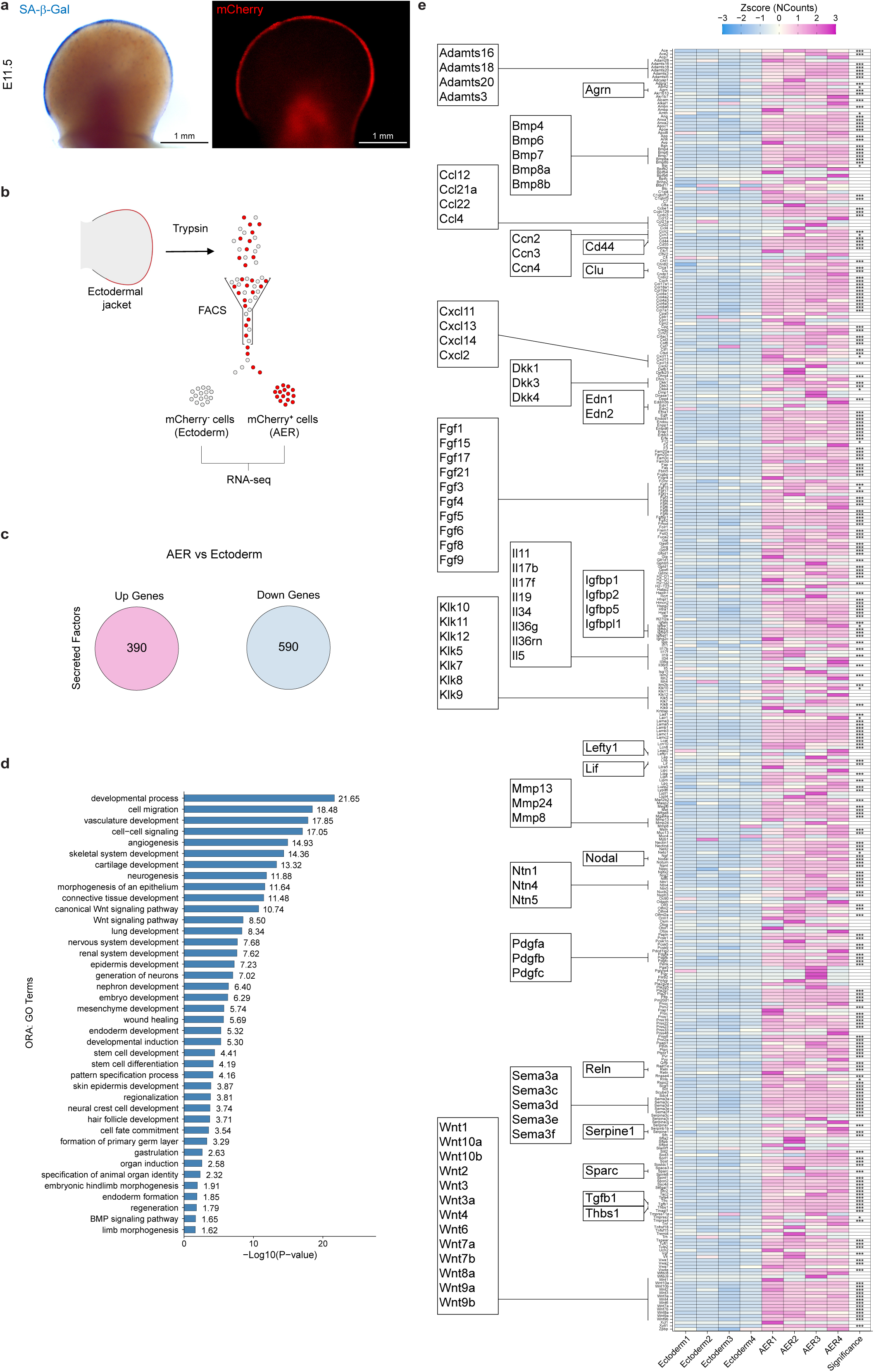
RNA sequencing of AER-derived p21+ senescent cells shows a significant developmental signaling potential. **(a)** SA-ß-Gal staining and mCherry visualization of embryonic day (E) 11.5 p21+/mCherry+ forelimbs. (**b**) Schematic of the strategy to isolate mCherry+ and mCherry-forelimb cells. Sorted mCherry+ cells, labeled as apical ectodermal ridge (AER), compared to mCherry-cells (labeled as ectoderm). (**c**) Number of genes encoding secreted factors exhibiting differential expression greater than 1.5-fold. (**d**) Select Gene Ontology (GO) terms from overrepresentation (ORA) analysis on upregulated secreted factors in c. (**e**) Heatmap representing normalized counts of the upregulated secreted factors in AER cells. Significant differentially expressed genes are labelled as follows: * (adjusted P < 0.05), ** (adjusted P < 0.01), and *** (adjusted P < 0.001).

Here, we wanted to explore the SASP of developmental senescence and ask if this is also conserved across senescence models. For this, we took the complete list of genes encoding “secreted proteins” from the Human Protein Atlas^21^, and overlapped this with the genes increased in the senescent AER, when compared to the non-senescent ectoderm. This identified 390 upregulated and 590 downregulated genes encoding secreted factors (Fig. 1c, Supplementary Table 1). As expected for cells originating from a developmental signaling center, over-representation analysis (ORA) of the upregulated genes showed several developmental pathways were overrepresented. These include terms related to development of multiple tissues and early embryonic processes like *neurogenesis, regionalization, gastrulation*, *primary germ layer formation*, along with *angiogenesis, wound healing*, *cell to cell signaling* and *cell migration* (Fig. 1d). In other contexts, the SASP is typically described as a mix of proteins including cytokines and inflammatory factors, but previous studies of developmental senescence using microarray analysis had noted an absence of these ^2,14^. The analysis here with RNA sequencing provides a more detailed developmental SASP profile. Focusing on members of the Ccl, Cxcl and Interleukin (Il) families, we found that the signature from developmental cells does not include classical SASP-associated cytokines including *Il1a*, *1b*, *6* or *Ccl2*, in agreement with previous studies. However, we identified that other molecules are expressed in developmental senescent cells, such as *Ccl4*, *12*, *22*, *Cxcl2*, *11*, *13*, *14* and *Il5*, *11*, *17*, 19 and others (Fig. 1e). We then looked more broadly at the genes encoding all secreted factors, to gain an unbiased profile of the SASP. Here, it was evident that many genes encoding growth factors, extracellular matrix and developmental signaling genes were increased in the senescent population. These included multiple members of the *Adamts, Bmp, Dkk, Fgf, Igfbp, Klk, Mmp, Ntn, Pdgf, Sema* and *Wnt* families, in addition to additional genes associated with early embryonic patterning and limb development, including *CD44, Agrn, Clu, Lefty1, Lif, Nodal* and others (Fig. 1e).

#### Developmental signaling is a common feature of in vitro senescence

As this analysis is the study of a major signaling centre that instructs tissue patterning and formation, it is expected that many developmental secreted factors are expressed. However, our interest was to determine if this signature is conserved in others models of senescence. To explore this, we first used publicly available RNA-seq datasets of cellular senescence in multiple contexts. We collected 18 datasets analyzing senescent cells from various mouse and human cell lines (IMR90, MEF, HAEC, HUVEC, Wi38, PMK, Pan02, HA549) that had been induced by different triggers such as Irradiation (IR), oncogene-expression (OIS), replicative exhaustion (RS), and chemotherapy (TIS) (Extended Data Fig. 1a). This allowed us to generate the differential expression profiles for all 18 senescence models compared to their respective controls (Fig. 2a).

**Figure 2.**
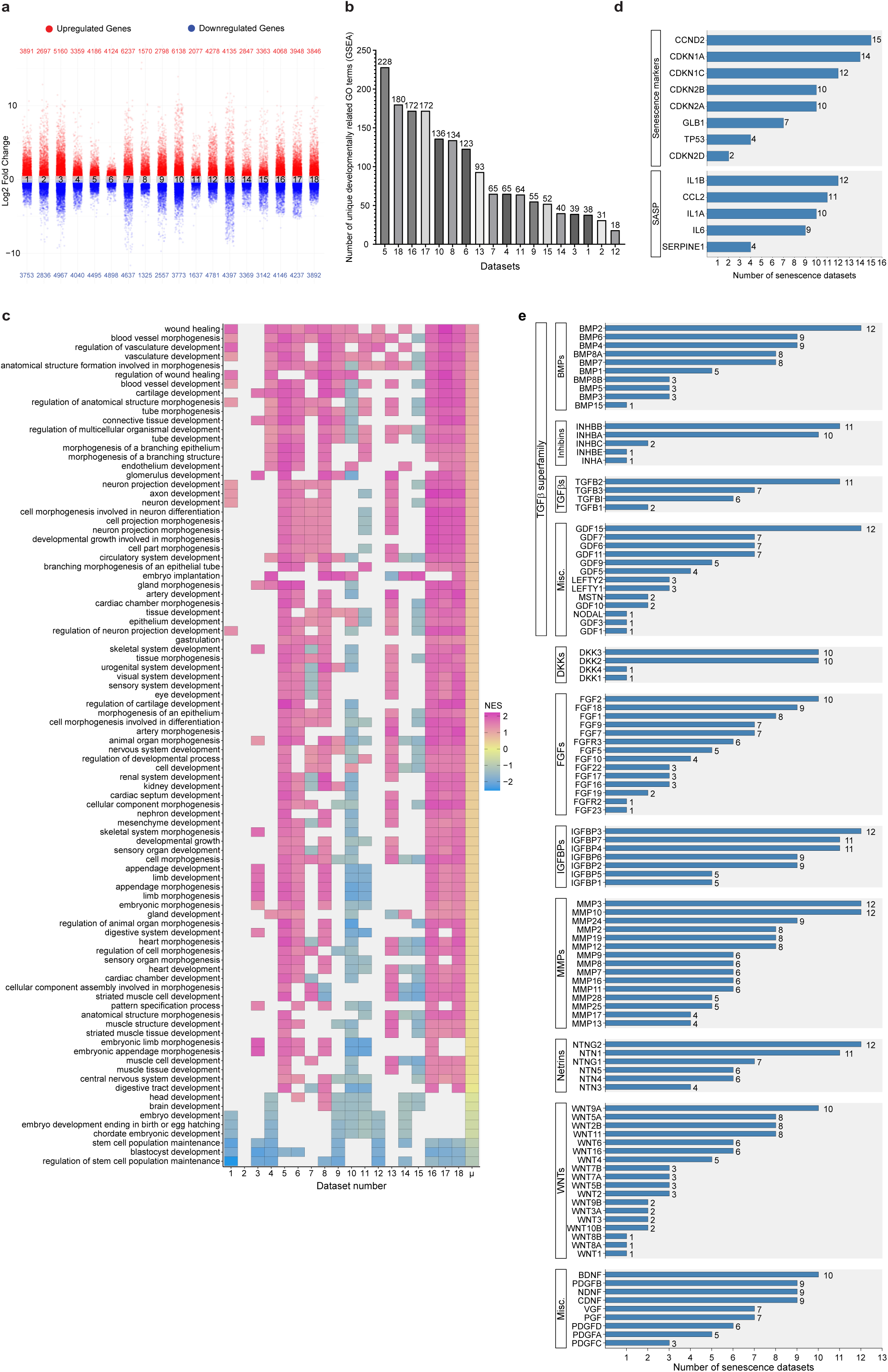
Transcriptome profiling of senescence models reveals signatures of processes and secretion linked to embryonic development. **(a)** Overview of the differentially expressed genes (DEG) across 18 different datasets comparing senescent cells to their proliferating controls. The DESeq2 workflow was performed to identify DEGs for each dataset separately. Genes averaging 25 counts were filtered out. Significant (adjusted P < 0.05) up- (red, log_2_ fold > 0) and down- (blue, log_2_ fold < 0) regulated genes in senescent cells were plotted along with the total significant gene count for each category. (**b**) Total number of significantly enriched Gene Ontology (GO) terms identified by GSEA for each dataset, that met our filter requirements for developmentally relevant biological processes. (**c**) Heatmap showing the identified developmental related GO terms if present in at least 7 distinct datasets. The X-axis shows the various datasets, with tiles showing their individual normalized enrichment score (NES) per term. ‘µ’ represents the average normalized enrichment score across every dataset and was used to sort the terms in a descending order. A positive NES value signifies that the given term is positively enriched in senescent cells. (**d**) Bar graph showing a subset of common senescence genes. X-axis represents the number of senescent datasets in which the selected genes are significantly upregulated (adjusted P < 0.05; >1.5-fold). (**e**) A selection of developmentally related genes coding for secreted proteins. The X-axis shows the number of datasets in which the selected genes are significantly upregulated (adjusted P <0.05; >1.5-fold change) in senescent cells compared to their proliferating controls.

First, we performed gene set enrichment analysis (GSEA) on each dataset. The top negatively enriched GO Terms shared between 17 or more senescence datasets were clearly related to senescence, mainly associating with mitosis and the cell cycle, supporting the analytic approach (Extended Data Fig. 1b). Then as we wanted to look for conserved developmental features, we used custom filtering criteria to isolate significant terms related to developmental-associated pathways. This led to the identification of 411 unique terms, with significant overlap across conditions (Fig. 2b). For instance, most datasets have more than 40 unique terms related to developmental processes. Irradiated HAEC cells have the most with 228 distinct GO terms (Dataset 5). In contrast, doxorubicin-treated pancreatic cancer cells (Pan02; dataset 12) only showed 18. To better compare the studies, developmentally related terms present in 7 or more senescence datasets were then represented graphically (Fig. 2c). From this, it is apparent that developmental processes are significantly and positively associated with senescent cells across all contexts, related to many diverse tissues, such as the development of the heart, vasculature, brain, limbs, and skeletal muscle. Additionally, there are more specific terms that were highly enriched, including *gastrulation*, *embryo implantation*, and *growth during development*. Interestingly, *wound healing* and *blood vessel morphogenesis* are among the most enriched terms shared by most datasets, which is in agreement with their previous association with senescent cells. Strikingly, datasets 16,17, and 18, which are A549 lung cancer cells in which senescence was induced by cisplatin, docetaxel, or palbociclib respectively, share the most developmental terms compared to the other datasets, suggesting there may be a link with TIS in cancer cells and developmental reactivation.

To further explore these patterns, over-representation analysis (ORA) was also conducted for each dataset, but using only significant genes (adjusted p-value < 0.05) that were up- and downregulated more than 1.5 log_2_ fold compared to their respective controls (Extended Data Fig. 1c). This uncovered largely similar results, and highlighted that the developmental signatures were associated with significantly upregulated genes in the senescent cells. Together, these results demonstrate that senescent cells, regardless of the mode of induction or cell type, show a strong association with developmental gene signatures. This association is driven by the upregulation of developmental genes, indicating that the senescent state entails a reactivation of developmental programs

Our main aim here was to ask if the SASP of these diverse senescence models shares any common signature with developmental senescence. As we wanted to generate a comprehensive picture of the SASP, one that was not just focused on inflammatory factors, we again used the complete list of genes encoding secreted proteins from the Human Protein Atlas, and overlapped this with the significantly upregulated genes with an expression level of more than 1.5-fold in senescent cells in each of the 18 senescence datasets. This resulted in a total of 937 genes encoding secreted proteins that are significantly upregulated in any of the datasets (Supplementary Table 2). It is known that the SASP is very heterogenous, and a primary determinant of SASP composition is the cell type that produces it^22^. However, by comparing the total list of genes for secreted proteins across all studies, interesting and common patterns emerged. It is clear that many typical SASP factors are frequently detected, including immuno-stimulatory factors, such as CCLs, CXCL, interleukins and others, as expected (Extended Data Fig. 1d). For example, Il1A, Il1B, and Il6, among the most frequently used SASP identifiers, are the most increased interleukins, along with Il33, in 9-12 different senescence datasets. For comparison, we also show the expression pattern for non-secreted senescence mediators, including p21 (*CDKN1A*) and p16 (*CDKN2A*), which shows that under similar parameters, these are also significantly increased in 14 and 10 datasets respectively (Fig. 2d).

However, what was striking from the list of secreted factors is that many genes that are linked to embryonic development are similarly upregulated in senescent cells, across all studies (Supplementary Table 2). This includes many members of the classical developmental pathways, including Fgf’s, Bmp’s, Wnt’s, Igfbp’s, and others (Fig. 2e). Interestingly, while there is heterogeneity in the expression of specific factors across all datasets, it is clear that each senescence condition activates members of key developmental families, while the individual factors might differ. For example, out of the 18 senescent contexts, *Fgf2* is significantly increased in 11 and *Fgf18* in 10, while *Bmp2* is increased in 12 and *Bmp8a* in 11. Similar patterns are seen for the larger Wnt and Tgf families, in addition to Igfbp’s and others, each of which play key roles in embryo development and organogenesis.

This analysis also shows that various other proteins that instruct development, and antagonists that contribute to fine-tuning the effects of these morphogens, are similarly increased (Fig. 2e). For example, the Growth Differentiation Factors (GDF), which are members of the TGF-β superfamily, are frequently increased in senescent cells. They regulate key aspects of embryonic axis formation, endoderm and mesoderm induction, kidney, pancreas or central nervous system (CNS) development^23^. In addition, Dickkopfs (Dkk) are extracellular modulators of Wnt signaling and play region-specific roles in establishing signaling gradients, with multiple roles in heart, limb, forebrain and muscle patterning^24^. Another relevant group are Netrins. These mostly secreted proteins, while traditionally found to be involved in axon guidance and development, they have also been shown to regulate organ morphogenesis during embryonic development via cell migration, adhesion and stem cell self-renewal^25^.

Some of these factors have already been associated with cellular senescence and the SASP in specific contexts, including some BMPs, Wnts, IGFBPs and others ^26–28^. Nevertheless, several of these secreted morphogens and growth factors implicated in instructing and regulating different embryonic tissues have not been characterized as integral components of the SASP in as extensive a manner. Collectively, these results imply that senescent cells acquire a strong developmental signaling potential as a core characteristic of the SASP.

### Senescent cells in muscle regeneration and aging display developmental signatures

This link with developmental signaling and senescence next led us to ask whether this is also observed in senescent cells in vivo. To address this, we used data generated from a recent study of muscle injury and aging^29^. Here, Moiseeva et al. provoked injury in muscle through cardiotoxin injection, inducing a transient wave of senescence that is prolonged in aged mice. Then, they sorted the senescent and non-senescent cells from the muscle, 3 (D3) and 7 (D7) days after injury, in both young and old animals, for RNA sequencing (Extended Data Fig. 2a). In this way, they generated a transcriptome map of senescence in vivo, in young and old satellite cells (SCs), fibroadipogenic progenitors (FAPs) and myeloid cells (MCs). We extracted the data for each condition, comparing the senescent to the non-senescent cells in each of these populations, in both young and old mice (Supplementary Table 3). We then performed GSEA analyses using identical metrics and thresholds as in the previous in vitro meta-analysis.

The total count of developmentally related GO terms was determined across all conditions, for each cell type (Extended Data Fig. 2b). We then displayed them by cell type in the young and old conditions, which revealed interesting dynamics (Fig. 3a-c). Indeed, there were many terms linked to development and the formation of muscle, but also blood vessel, brain, heart, lung, kidney and many other tissues. And again, terms related to early development including *gastrulation*, *germ layer formation* and *patterning* were prominent, with interesting dynamics across the stages. For example, satellite cells at D3 and D7 share a common trend of associated developmental terms in young mice (Fig. 3a). However, in old mice, there are now many more terms at D3 after wounding, but which are not maintained by D7. In FAPs, both young and old cells at D3 share a similar pattern, but, while this decreases somewhat by D7 in young mice, it is completely absent in aged mice, suggesting this property is not properly maintained (Fig. 3b). Curiously, young myeloid cells have a negative association with developmental terms, but which changes in aged animals, where there is a pronounced positive enrichment of these properties at D3 (Fig. 3c). Collectively, the data indicates various points. First, that senescent cells in vivo in these three contexts have a pronounced association with developmental signatures when compared to their non-senescent same-aged counterparts. Second, these developmental signatures appear to be altered in aging mice.

**Figure 3.**
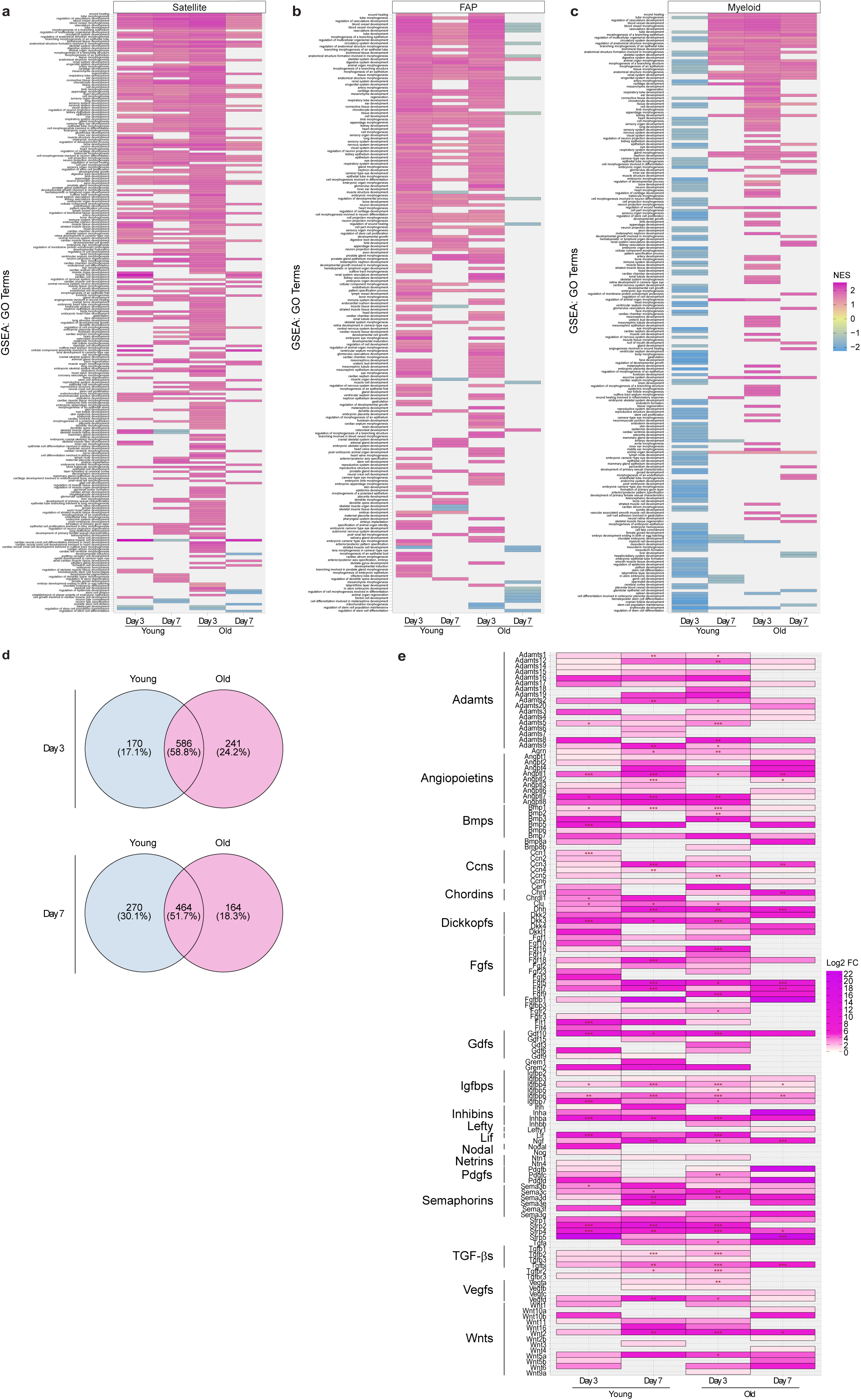
Senescent cells display increased transcriptional profiles related to developmental programs in both young and aged murine skeletal muscles. (**a, b, c**) GSEA was performed on bulk RNA sequencing data from Moiseeva et al., from (a) Satellite, (b) Fibro-adipogenic progenitor (FAP) and (c) Myeloid cells, three- or seven-days post cardiotoxin injection, from young (3-6 months) or old (28 months) mouse muscles. For each condition senescent cells were compared to their non-senescent counterparts and GO terms related to developmental biological processes were identified. Of these, the 60 most positively enriched terms according to the normalized enrichment score (NES) across all comparisons were selected and plotted separately for each cell type. Y-axis shows one distinct term per line. A positive NES value signifies the term is positively enriched in senescent cells. (**d**) Venn diagram showing the total number of upregulated genes (>1.5-fold) coding for secreted proteins in senescent satellite cells from young and old mice, 3- or 7-days post cardiotoxin injection. Numbers refer to total gene counts or percentages of the total. (**e**) Selected genes for secreted factors were plotted for each condition comparing senescent to non-senescent cells. Significant differently expressed genes are labelled as follows: * (adjusted P < 0.05), ** (adjusted P < 0.01), and *** (adjusted P < 0.001).

Focusing on the satellite cells, we next examined the SASP, looking again at the full complement of secreted factors. We extracted genes encoding secreted factors that were more than 1.5-fold upregulated in senescent cells compared to their non-senescent controls (Fig 3d, Supplementary Table 4). Although both young and old senescent satellite cells maintain a similar number of shared genes at three- (586 genes, 58.8%) and seven- (464 genes, 51.7%) days post-injury, aged senescent cells have an increased proportion of exclusive genes at D3 compared to young (24.2% vs. 17.1%), but which is then decreased by D7 (18.3% vs. 30.1%). This suggests that while the core number of secreted factors may remain constant over time, the secretory signature of senescent cells becomes altered with age.

We then focused on the expression of all of the upregulated genes for secreted factors at each of the 4 timepoints (Supplementary Table 4). Looking first at known SASP factors including cytokines, chemokines and interleukins, there is a clear upregulation of many of these in the senescent cells compared to their non-senescent counterparts, and with interesting dynamics across the different timepoints (Extended Data Fig. 2c). For example, IL6, a hallmark SASP factor is increased in young SCs at D3 and D7 after injury, but while also induced in old SCs at D3, expression is not maintained at D7. Similarly, Ccl2, another common SASP factor, is also significantly up at all timepoints, except in old cells at D7. When we looked at the list of developmentally-associated factors, again, it was clear that there is a pronounced induction of these factors in senescent cells compared to their non-senescent equivalents (Fig. 3e for selected factors; Extended Data Fig. 3d for extended heatmap). Many belonged to families previously identified in vitro, including *Bmp’s*, *Fgf’s*, *Gdf’s*, Igfbp’s, Wnt’s, and others. Interestingly, the expression dynamics of these factors differed between young and old tissue. Several signaling molecules—such as *Bmp7*, *Dhh*, *Dkk2*, *Fgf5/7/18*, *Gdf10*, *Igfbp4/6/7*, *Inhba*, *Ngf*, *Sfrp1/4*, and *Wnt2/5a*—were consistently upregulated in senescent cells across all time points and age groups. However, a subset of factors, including *Bmp6*, *Fgf3/10*, *Gdf9*, *Nodal, Ntn4* and *Sema3f*, were specifically enriched in young senescent cells, while conversely, other factors including *Bmp8b, Fgf17/9, Gdf3, Igfbp5, Inhbb, Lefty1, Tgfb1, Vegfc, Wnt10a/2b/4/9a* were uniquely induced in old senescent cells. It is also particularly curious to note again that several genes associated with gastrulation and early embryonic patterning—such as *Inhba*, *Cer1*, *Chrd*, *Dkk1*, *Lefty1*, *Nodal*, *Nog*, and *Wnt3*—were induced in senescent cells at distinct stages. Overall, however, this analysis of the full complement of SASP factors from senescent cells in young and old muscle supports that senescent SCs do indeed reactivate developmental instruction mechanisms, but that this may change with age.

### The SASP can activate developmental-fate genes in a dose-dependent manner

Next, we wanted to functionally test if senescent cells have the potential to instruct developmental fate. We had previously shown that the SASP can induce dedifferentiation and cell plasticity, using oncogene- and irradiation-induced senescence in primary mouse keratinocytes (PMK) (a model similar to dataset number 11 in Fig. 2a)^13^. In particular, we found that transient 2-day exposure to the SASP caused an upregulation of skin stem cell genes, and that the cells receiving this stimulus reprogrammed into functional hair follicle stem cells. However, prolonged SASP exposure for 6 days led to increased expression of skin stem cell genes, but also caused paracrine senescence, presumably as an intrinsic barrier to increased plasticity.

Following from this, we were interested to explore the extent of SASP-mediated reprogramming, and performed bulk-RNA sequencing on PMK exposed to the SASP from oncogene-induced senescence (OIS) in PMK, for either 2- or 6-days (Fig. 4a, Supplementary Table 5). After 2 days of SASP exposure, there were 615 significantly differentially expressed genes (DEGs), with 334 upregulated and 113 downregulated genes. However, 6 days of exposure to SASP, which led to paracrine senescence, resulted in the differential expression of 7,922 genes in PMK, including 3511 upregulated and 1,684 downregulated more than 1.5-fold (Fig. 4b).

**Figure 4.**
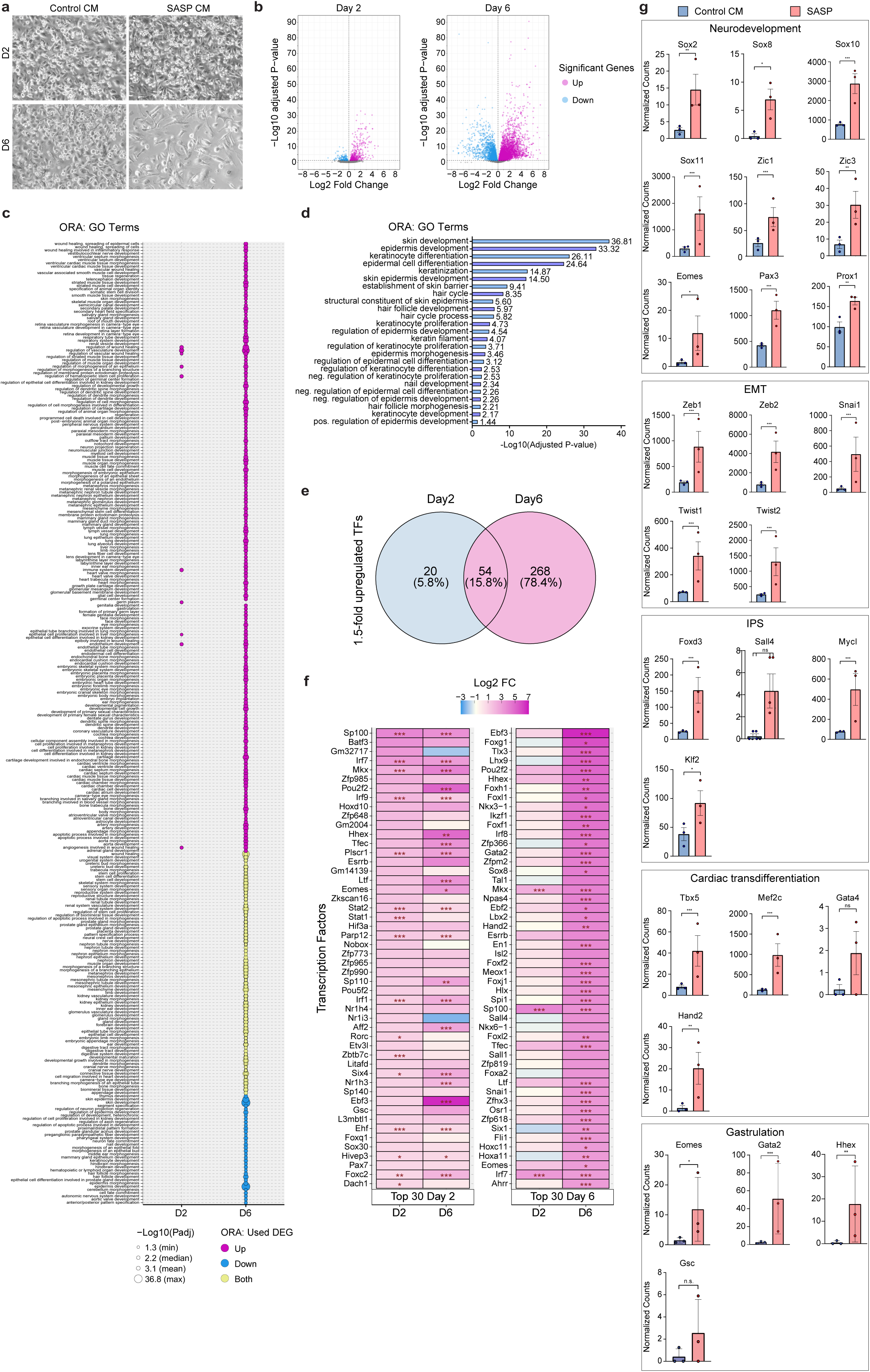
SASP exposure induces developmental gene expression in primary mouse keratinocytes. **(a)** Phase-contrast images of primary mouse keratinocytes (PMK) 2- or 6-days after treatment with conditioned media (CM) obtained from either: Puromycin IRES GFP vector- (PiG, Control CM) or Harvey Rat sarcoma virus- (HRAS, SASP CM) plasmid transfected cells. **(b)** RNA sequencing was performed on PMK (n=3) treated with CM or SASP for 2x24h (Day 2) or 6x24h (Day 6). Volcano plot showing differentially expressed genes (DEGs) comparing SASP treated cells against PiG CM treated controls. Genes averaging less than 25 counts were excluded. Significant DEG (adjusted P < 0.05, and >1.5 or < -1.5-fold) are highlighted in color. **(c)** Overrepresentation analysis (ORA) was performed using the Gene Ontology (GO) database for significantly (adjusted P < 0.05) upregulated (>1.5-fold) and downregulated (< -1.5-fold) genes following 2x24h (D2) and 6x24h (D6) PMK treatment with CM/SASP. Overrepresented GO terms associated with developmental processes were selected and visualized; upregulated genes in magenta, downregulated in blue, and yellow if a term was linked to both. **(d)** Skin related GO terms overrepresented among significantly downregulated genes in PMK treated for 6x24h with SASP. For both (c) and (d) P-value was determined significant if adjusted P < 0.05. **(e)** Venn diagram illustrating upregulated transcription factors (TF) (>1.5 fold) in PMK upon SASP treatment for either 2 or 6 days. Data is presented as total TF counts or percentages of the total. **(f)** Heatmap with two separate panels, each showing the top 30 upregulated TFs following 2 days (left) or 6 days (right) SASP treatment, relative to control CM. For each panel, TFs were ranked by decreasing log_2_ fold change relative to the panels respective day and expression is shown for both treatment durations. Significantly upregulated genes are indicated with * (adjusted P < 0.05), ** (adjusted P < 0.01), and *** (adjusted P < 0.001). **(g)** Total normalized counts of several TFs associated with distinct biological processes for neurodevelopment, epithelial to mesenchymal transition (EMT), induced pluripotent stem cells (IPS), cardiac transdifferentiation and gastrulation, in PMK exposed to SASP and control CM following 6 days of exposure. Significant differently expressed genes are marked as follows: * (adjusted P < 0.05), ** (adjusted P < 0.01), and *** (adjusted P < 0.001).

To further explore the DEGs, we performed GSEA analysis on both timepoints. As expected, the top signatures reflected the SASP influence on the cells at both D2 and D6, and included many signature terms related to inflammation, interferons, damage response and senescence (Extended Data Fig. 3a). However, in line with our hypothesis, we noticed a pronounced association of terms related to embryonic development. Indeed, when we selected for terms associated to development, stem cells and regeneration, there was a striking association in the upregulated genes in the SASP treated cells, particularly after prolonged SASP treatment (Extended Data Fig. 3b). Interestingly, most of the few terms associated with 2-day SASP exposure related to blood vessel and wound healing, known features promoted by the SASP. But then, with longer SASP exposure, additional terms related to many developmental processes appeared, including those relating to stem cell development, neuron development, and formation of various tissues including kidney, heart, lung, eye and others. To explore this further, we used ORA analysis to look at all of the significantly different genes with >1.5-fold change, which again showed a striking association with developmental fate in the 6-day SASP-exposed cells (Fig. 4c). In fact, developmental terms associated with uniquely upregulated genes represent 10 terms following 2 day-SASP exposure, but 188 following 6 days of SASP treatment. Focusing on the 6-day SASP exposure, it is striking that the top developmental terms relate to embryonic development of multiple tissues, including blood vessel, muscle, nervous system, kidney, lung and many more. And again, there is a strong association with very early embryonic development, including *gastrulation*, *formation of primary germ layer*, *paraxial mesoderm development* and others. These results uncover that the SASP can indeed activate genes not only associated with angiogenesis and wound repair, but also that prolonged SASP exposure leads to increased expression of genes associated with very early development, and of multiple tissues.

Conversely, when looking at the ORA results for the most significantly changing genes, downregulated genes do not overrepresent developmental terms after a short SASP treatment, and only represent 34 terms after prolonged SASP exposure (Fig. 4c). Interestingly, the signatures associated with the downregulated genes largely relate to skin signatures, ranging from *keratinocyte* and *epidermis differentiation* to *hair follicle* and *keratinocyte-development*, suggesting a repression or loss of original cell identity (Fig. 4d). Overall, this suggests that chronic SASP exposure can alter cell fate, in this case, decreasing skin and ectodermal fate signatures (in keratinocytes), and increasing diverse early developmental signatures of other tissues and germ layers.

A central tenet of embryonic development is the expression of key transcription factors (TFs) that define and instruct key cell populations and fates. To investigate if developmental TFs are induced by the SASP and associated with the developmental signatures, we extracted a complete list of TFs that were increased in the 2-day and 6-day SASP-treated cells. This turned out to be quite extensive, with 74 TFs increased by >1.5-fold in response to 2-day SASP, and 322 in response to 6-day SASP (Fig. 4e, Supplementary Table 6).

In agreement with the previous pathway analysis, we found that many TFs that are known to play key roles in various stages of development were increased by the SASP, and that this effect appears to be dose responsive, with longer exposure activating more genes (Fig. 4e). For example, when we looked at the top 50 TFs induced by two-day SASP exposure, this included genes associated with early development and reprogramming such as *Hhex*, *Esrrb*, and *Eomes*, which were increased further with six days exposure. However, many additional TFs were then appreciably induced with prolonged SASP treatment, such as *Foxg1*, *Foxl1*, *Gata2*, *Meox1* and *Sall4*. Initially being undetected or having low expression levels after two days of SASP treatment, these TFs all increased after prolonged exposure (Fig. 4g). Looking at further associations with developmental fate, it was clear that six-day exposure to the SASP induced expression of TFs associated with many and distinct developmental processes: for example, genes associated with neurodevelopment (incl. *Sox2*, *Sox8*, *Sox10*, *Sox11*), epithelial-mesenchymal transition (EMT) (*Snai1*, *Twist1*, *Twist2*, *Zeb1*), iPS reprogramming (*Foxd3*, *Mycl*, *Sall4, KLF2*), cardiac trans-differentiation (*Mef2c*, *Hand2*, *Tbx5*, *Gata4*) and Gastrulation (*Foxh1*, *Gata2*, *Eomes*, *Gsc*) were all increased. Altogether, these results uncover an unexpected and previously undescribed capacity of the SASP to induce expression of a diverse set of developmental genes, including key TFs, that are associated with many stages of embryonic development.

### Therapy-induced SASP induces phenotypic changes and developmental genes in cancer cells

Finally, we wanted to explore our hypothesis using an independent senescence model to those already examined, but which might not undergo paracrine senescence in response to prolonged SASP. In a proof-of-concept experiment, we used doxorubicin to induce senescence in the liver carcinoma cell line Huh7, and examined this by bulk RNA sequencing for developmental SASP factor-expression. Then, we treated proliferating Huh7 cells with the SASP from these cells for 6 days, and checked for developmental gene induction, again by RNA sequencing (Fig. 5a).

**Figure 5.**
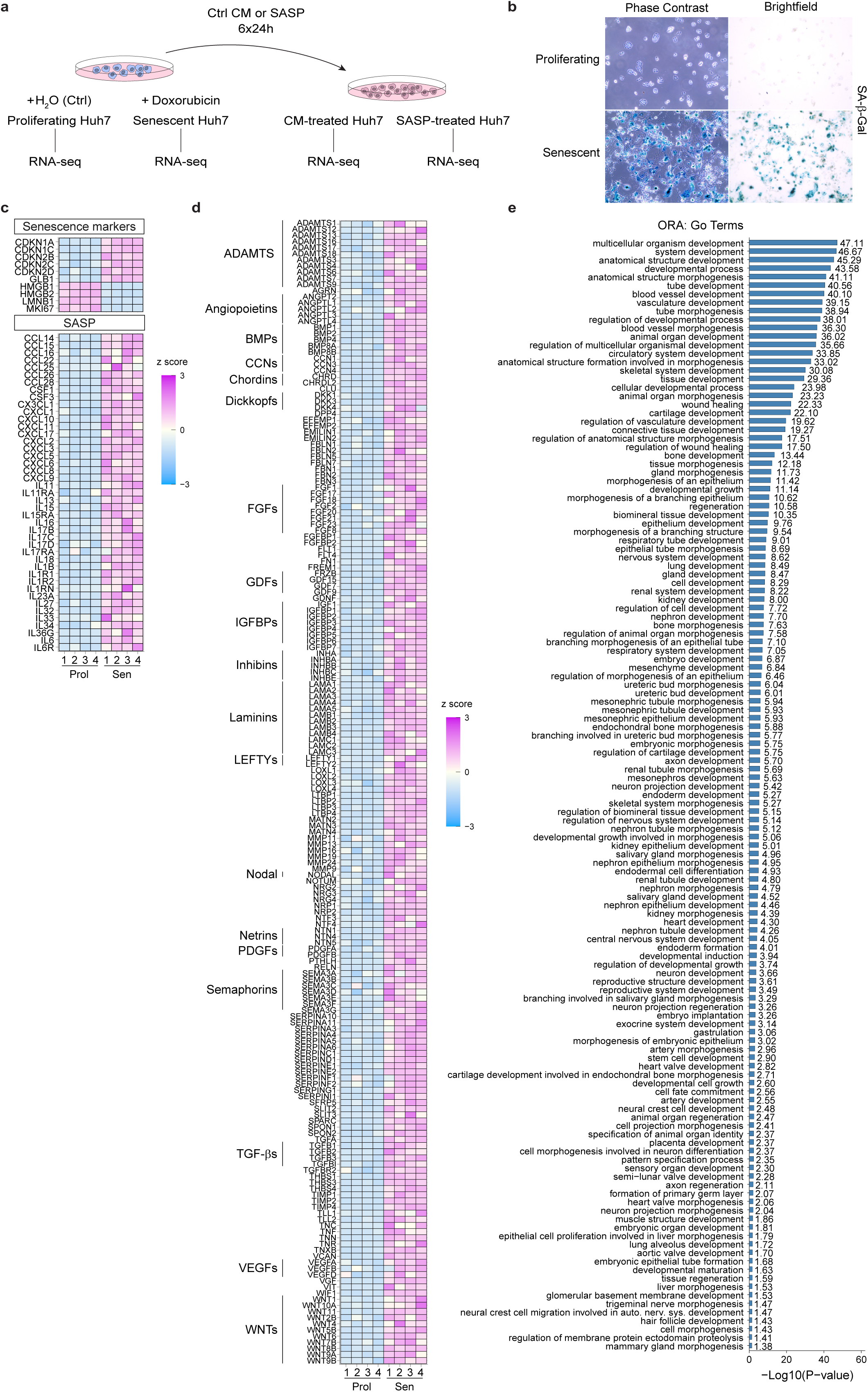
Therapy-induced senescence in Huh7 liver cancer cells is associated with diverse developmental signaling signatures. **(a)** Schematic representation of the strategy used to generate proliferating and senescent cells, and control conditioned media- (CM) or SASP-treated Huh7 cells, for RNA sequencing analysis. Senescence was induced using 200nM of doxorubicin and the SASP was collected 10 days after induction began. The SASP, or control conditioned media (CM) from proliferating cells, was used to treat Huh7 cells for 6 days. RNA from proliferating and senescent, and CM- and SASP-treated Huh7 cells was used to perform bulk RNA sequencing. (n=4 each) (**b**) Phase Contrast and Brightfield images of proliferating and senescent cells, stained for SA-ß-Gal activity (**c**) Heatmap showing z-score adjusted normalized counts for a select set of senescence-and SASP-associated genes (adjusted P < 0.05) from bulk RNA sequencing of proliferating (Prol) and senescent Huh7 cells (Sen) n=4. (**d**) Similar heatmap as in (c) but showing a representative selection of genes known to be secreted and involved in developmental processes (adjusted P < 0.05; upregulated >1.5-fold). (**e**) Overrepresentation analysis (ORA) GO terms using genes coding for secreted factors in senescent Huh7 cells (adjusted P < 0.05; >1.5-fold.)

First, we treated Huh7 cells with doxorubicin to induce senescence. Seven days after treatment, the cells exhibited classical features of senescence, including enlarged size, no proliferation and high SA-β-Gal activity (Fig. 5b). These samples were then analyzed by bulk RNA sequencing, which further confirmed the senescent state, revealing elevated expression of key senescence genes including increased p21 (*CDKN1A)* and decreased *LMNB1* (Fig. 5c: Supplementary table 7). In addition, several genes associated with the SASP, including various cytokines and inflammatory factors, were also significantly upregulated.

As before, we next extracted the list of genes encoding secreted factors, and analyzed those which were significantly increased compared to proliferating controls (Supplementary Table 8). In agreement with our hypothesis, the same developmental families including *BMPs, FGFs, IGFBPs, WNTs* and others were highly enriched (Fig. 5d). In particular, there is robust activation of many members of the WNT family, such as *WNT6*, *10A*, *5B* and others, or the FGF family, including *FGF21*, *17*, and *8*. And again, there was notable induced expression, although at low levels, of early embryo and gastrulation-associated factors, including *CHRD*, *DKK1*, *INHBA*, *LEFTY1*, *2*, and *NODAL*. These results confirm that senescent cells activate or increase expression of many developmental signaling factors.

Next, we performed pathway analysis on this list of secreted factors that were increased in senescent cells, selecting as before for developmentally-related terms (Fig. 5d, Supplementary Table 9). This produced a striking association that was increased in the senescent state. Again, these terms did not just relate to the cell-type of origin but reflected broader developmental programs, including *wound healing* and *blood vessel development*, but also *endoderm formation*, *gastrulation* and *neuron development,* as well as development of many tissues including lung, kidney, heart, bone and others.

Finally, we asked if this chemotherapy-induced SASP could induce developmentally-associated genes in cells exposed to the SASP. For this, we did as we had with mouse keratinocytes, and exposed Huh7 cells to the SASP for 6 days, before analyzing by bulk RNA sequencing (Supplementary Table 10). Interestingly here, repeated exposure of Huh7 cells to the SASP did not induce paracrine senescence. In agreement with previous findings, the SASP actually increased colony size, but with decreased layering of colonies, and increased cellular heterogeneity and migration (Fig. 6a). Pathway analysis on these cells yielded fewer developmental terms than in the PMK, but did again associate with blood vessel development and wound healing as in the other contexts. This also revealed broader developmental terms including *heart field specification*, *tube morphogenesis*, and *regulation of developmental growth* (Fig. 6b). To look for activation of developmental genes, we again focused on TFs that were significantly induced by the SASP in these cells. While this was also not as robust as with PMK, possibly related to the continued proliferation of the SASP-treated cells, we did identify similar trends, with increased expression of genes related to those seen in PMK, including *KLF4*, *6*, *GATA2*, *MEF2C*, *HAND1*, *TBX2*, *4*, Egr1, numerous Hox genes including *HOXA1*, *13*, *HOXB3*, *9* and others (Fig. 6c, Supplementary Table 11). Overall, these experiments provide functional evidence that senescence induction involves reactivation of developmental signaling properties, that can induce expression of key developmental fate genes.

**Figure 6.**
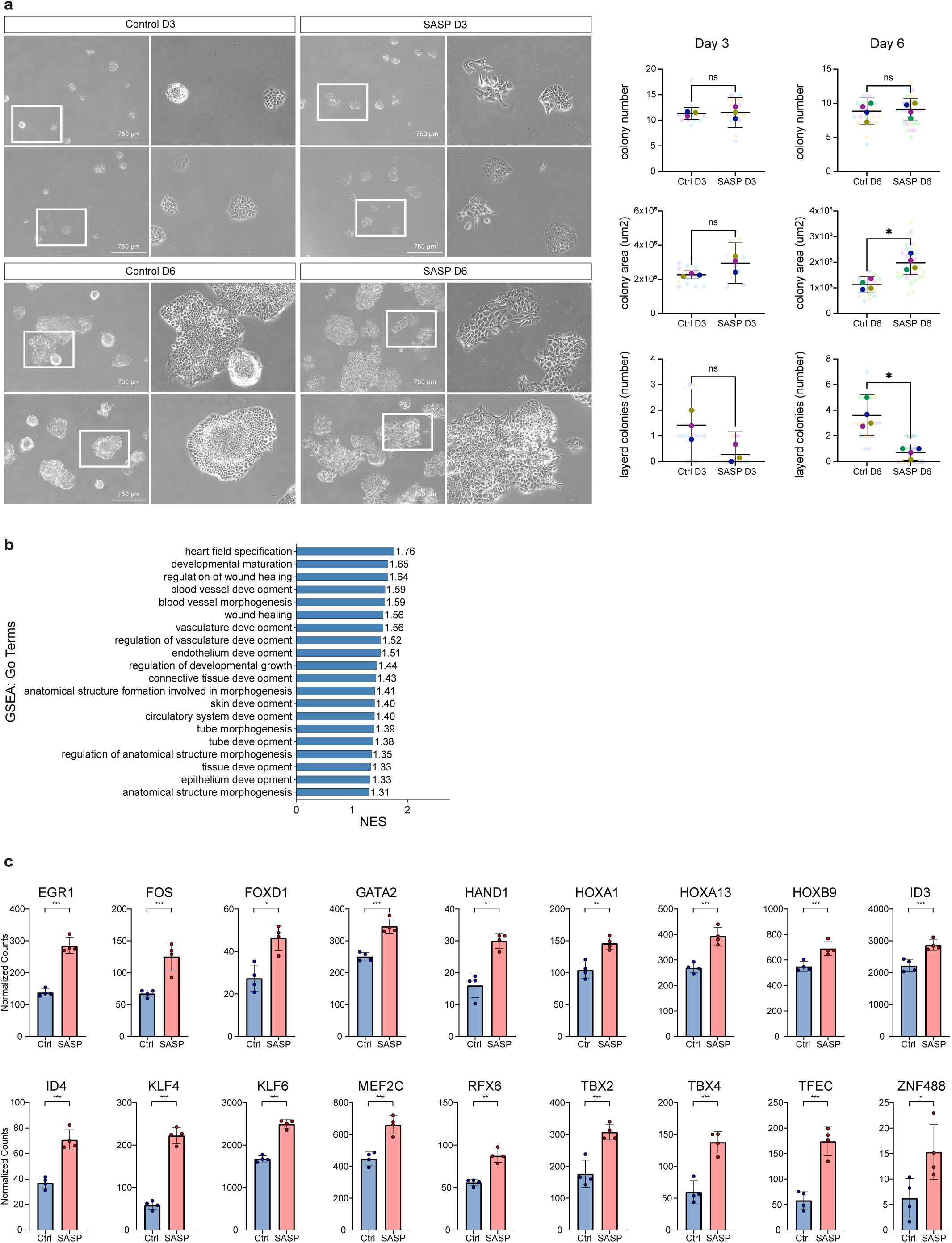
SASP treatment induces phenotypic and transcriptional changes in Huh7 cells. **(a)** Phase Contrast images of Huh7 cells treated for 3 or 6 days with control conditioned media or SASP. For each condition, two example images are shown. Images on the right are digital zooms of the area outlined by a white rectangle. Superplots show measurements of colony numbers, colony area and the number of multilayered colonies. An unpaired Mann-Whitney test was used to test statistical significance (*; P < 0.05, ns; not significant). (**b**) Developmentally relevant and significantly enriched terms from GSEA using GO terms, comparing SASP- versus CM-treated Huh7 cells. (**c**) Expression values of a selection of transcription factors upregulated by the SASP in Huh7 cells (SASP) compared to control CM treated cells (Ctrl).

## DISCUSSION

Here we identify that senescent cells across many contexts exhibit features and properties of developmental signaling centres, and make a provocative claim that reactivation of this developmental instructive state is a core feature of senescent cells. This analysis adds significantly to our conceptual understanding of senescence, with far-reaching implications for how its roles in health, development, and disease are interpreted and potentially manipulated.

Developmental signaling centers are defined as transient populations of cells within the embryo that secrete morphogens, such as WNTs, FGFs, BMPs, and SHH to direct cell fate, proliferation, and tissue patterning in surrounding cells ^17–19^. Indeed, such functional properties could already be ascribed to senescent cells in many contexts. Classical signaling centers such as the node, the apical ectodermal ridge (AER), the hindbrain roof plate (HRP), and the zone of polarizing activity (ZPA) operate via precisely such mechanisms, establishing spatial gradients of signals that instruct complex tissue morphogenesis. For example, the AER produces a gradient of FGF and WNT signals that instruct limb development and patterning^17^.

Previously, we showed that the AER exhibits classical features of senescence such as p21 and p15 expression, low proliferation, high secretion and SA-β-Gal activity^14^. Importantly, these markers are not restricted to a subset of cells within the AER; rather, the entire structure exhibits a uniform senescence-like profile. This raised an intriguing question: does the AER stain positive for senescence markers because it is functionally senescent, or do senescent cells across various contexts activate these markers because they are re-engaging a developmental signaling program? This question challenges conventional definitions of senescence, suggesting that these two phenomena may be more deeply intertwined than previously recognized.

This study builds on our concurrent study using a p21-mCherry-CreERT2 reporter mouse model to isolate and profile the AER^20^. We showed that senescent cells and the AER share a core gene expression signature of genes encoding non-secreted proteins, including *Eda2r*, supporting the idea that senescent cells may recapitulate features of developmental signaling centres. In this current study, we focus on the secretory profile of the AER and find that it includes, as expected, a broad range of classical developmental signaling molecules, such as BMP’s, FGF’s, IGFBP’s, and WNT’s. Remarkably however, by comparative analysis, we show many of the same factors expressed in the senescent AER are also consistently upregulated across diverse senescence models. Indeed, a transcriptome study of eighteen in vitro senescence datasets reveals that these developmental factors are as frequently induced as canonical SASP components like *Il1a*, *Il1b*, and *Il6*, suggesting that family-level activation of morphogen signaling is a conserved feature of the senescence program. As a major challenge in the senescence field is a lack of markers identifying senescent cells, the identification here that they also exhibit signaling overlap, when compared to their non-senescent counterparts, may further help with the identification of these cells, and provide additional SASP factors that can be tested for validation. Indeed, factors such as BMP2, DKK2, FGF2, IGFBP3, INHBB, NTN1, SRPX2, TNFSF15, or WNT9A, which are some of those significantly increased in more than ten out of eighteen senescence studies, may serve as potential SASP markers. Although this analysis does not assess expression levels per se, the repeated involvement of these signaling families points to a structured, conserved SASP that extends beyond inflammatory signaling. Of course, the genes identified here are not restricted to functioning only in developmental contexts, and many also play critical roles in adult tissue homeostasis and disease. We emphasize their developmental functions to underscore the observation that senescent cells share features with developmental signaling centers, thereby providing a framework to reconsider how senescence may contribute to tissue biology.

Importantly, this conservation is observed not only in cultured cells, but also in vivo in regenerative and aging contexts such as seen in muscle after damage, suggesting that senescent cells across diverse physiological and pathological settings engage developmental signaling modules. What is strongly supportive of this hypothesis is that in the muscle, gene-expression comparisons are made within the same cell type and age, directly contrasting senescent and non-senescent subpopulations (e.g., SA-β-Gal–positive versus SA-β-Gal– negative satellite cells) at each timepoint. This unequivocally demonstrates the increased expression of these developmental signaling factors is coupled with senescence markers. Interestingly, senescent satellite cells from young tissue robustly activate these pathways, whereas those from aged tissue show a dampened or dysregulated response. This is also evident for FAPs. Conversely, myeloid cells seem to acquire a developmental-like fate only in aging. This supports that in vivo, senescent cells also utilize developmental instructive mechanisms, but that age-related alterations may impact the fidelity of this developmental reactivation.

Critically, using two separate models, we demonstrate that the developmental signaling activity associated with senescent cells and the SASP is sufficient to induce the expression of developmental genes in other cells. This provides functional evidence that senescent cells can act as signaling centers, capable of influencing cell fate and identity through paracrine mechanisms. While senescent cells are known to accumulate during aging and in various diseases, the precise ways in which they impact surrounding tissue remain confusing. Notably, recent studies have linked aging and disease with the reactivation of developmental pathways and the emergence of alternate or aberrant lineage programs ^30–33^, yet the cellular sources and mechanisms driving these changes are often unclear. Our findings suggest that senescent cells may play a direct role in this process, by re-initiating developmental signaling cascades. This opens new avenues for investigating how senescence contributes to age-related dysfunction and offers potential strategies to either mitigate or harness these effects therapeutically.

Phenotypic plasticity is also a hallmark of cancer, associated with loss of markers of the cell of origin, or increased expression of genes associated with other tissues^32–36^. In addition, developmental- or onco-fetal-reprogramming is increasingly shown to contribute to tumor progression and recurrence^37–40^. As senescent cells have an intricate relationship with cancer, being present at all stages including in pre-malignant lesions, the tumor microenvironment, and following therapy^41^, the finding that they can induce developmental or lineage-inappropriate gene expression in other cells could open new avenues to disrupt cancer progression and recurrence. For example, we show that OIS in mouse keratinocytes, an in vitro model of papilloma, leads to activation via the SASP of a wide variety of stem and developmental genes, many of which have pro-tumorigenic roles, and in a dose responsive manner. We also demonstrate that therapy-induced senescence (TIS) in liver cancer cells can, through the SASP, upregulates genes associated with developmental fate, including *KLF4, MEF2C, GATA2, HAND1*, and multiple *HOX* family members. Notably, many of the transcription factors induced by the SASP are known drivers of tumor growth, heterogeneity, and metastasis. Further, the recurrent association between senescence and genes normally activated during gastrulation is also unexpected and intriguing. Gastrulation represents a critical developmental stage during which germ layer identity is specified, and the reactivation of genes linked with this program in senescent and SASP-treated cells— particularly in the context of cancer and TIS—warrants detailed investigation.

Overall, our discovery that senescent cells exhibit molecular and functional features of developmental signaling centers redefines the conventional view of cellular senescence as solely a terminal, damage-induced state. Instead, it suggests a broader paradigm in which senescence may represent a conserved, instructive cell state capable of orchestrating tissue architecture and intercellular communication, reminiscent of embryonic patterning hubs, and points towards new ways in which this process may be therapeutically targeted.

## MATERIALS AND METHODS

### Animals

C57Bl/6J (WT), and p21-mCherry-CreERT2^20^ mice strains were housed in a temperature- and humidity-controlled environment at the Institut Clinique de la Souris (ICS) animal facility of the IGBMC. The experimental procedures were in full compliance with the institutional guidelines of the accredited IGBMC/ICS animal house. They complied to both French and European Union regulations governing laboratory animal research, and were authorized by the French Ministry of Agriculture and Fisheries for animal experimentation.

### Isolation and culture of Primary Mouse Keratinocytes

Primary mouse keratinocytes (PMK) were obtained from the skin of C57Bl/6J newborn mice, no older than 24 hours. The euthanasia procedure comprised of one injection per animal (5µL ketamine and 2.5µL xylazine in 12.5µL NaCl 0.9% saline solution) administered intraperitoneally. The skin samples were dissected, washed with PBS and placed epidermis up on PBS with 2.5 mg/mL dispase II, overnight (Sigma 4942078001) at 4°C. The next morning, the dermis was separated from the epidermis. The epidermis was first minced and incubated in DMEM (4,5g/l glucose, Gibco™ 41965) supplemented with 10% FCS (Fetal Calf Serum) and 100 U/ml penicillin-streptomycin for 30 minutes at 37°C and subsequently filtered through a 40µm cell strainer. Isolated cells were plated and subsequently cultured at an initial density of 1 skin for a 20cm^2^ surface in collagen coated 6-well plates (Corning® BioCoat™ Collagen IV) in keratinocyte media, which included 450 ml EMEM, 40 ml chelated FBS (8%), 5 ml penicillin/streptomycin (1%), 250ul EGF 20ug/ml (10 ng/ml), 11,25ul CaCl2 2M (0,05mM). Isolated PMK were cultured at 37°C in 5% CO^2^.

### Viral transduction

Retrovirus was produced by transiently transfecting the Phoenix packaging cell line (G. Nolan, Stanford University, Stanford, CA) with helper virus (gift from Dr. Scott Lowe), and MSCV PIG plasmid (Scott Lowe, Addgene plasmid #18751) or p-Babe HRas^V12^ (Bob Weinberg, Addgene plasmid #1768). Viral vectors were harvested 48h post transfection and the supernatant was filtered with a 0.45 µm PVDF filter (Merck Millipore, SLHV033RB) and used to infect PMK for 2 hours first before repeating it for 24h and washing the cells with PBS. 48h post infection cells were split 1:2 into puromycin selection (1 µg/mL) for another 48h.

### Conditioned media and SASP collection

Conditioned media (CM)/SASP was collected from PiG or Hras^V12^-infected PMK 7 days after infection, and was filtered through a PVDF 0.45µm syringe filter. Collected CM was stored at -20°C until subsequent treatment of newly isolated PMK, every 24h for up to 6 days. Regarding Huh7 cells, (donated by T. Baumert, University of Strasbourg), these were grown in DMEM (4,5g/L glucose (Gibco™ 11965092) supplemented with 10% FCS, 1mM PyrNa, non-essential amino acids (Gibco™11140050), and 40ug/mL gentamycin). For senescence induction and SASP production, 1.75 × 10^6 Huh7 cells were seeded in P10 dishes and treated with 200nM of doxorubicin (44583-1MG, Sigma) for 96h. Doxorubicin treatment was repeated after the first 48h. Cells were then washed twice and cultured while changing media every 2 days. On day 6, medium was replaced and collected 24 h later. For control CM, due to the proliferative nature of the cells, an initial seeding density of 0.328 × 10^6^ cells was used and treated with H_2_O instead of doxorubicin. Medium was replaced on day 2 and collected 24h later. Both, CM and SASP were filtered with a PVDF 0.45µm syringe filters (Merck Millipore, SLHV033RB), were aliquoted and stored at -80°C prior to use. Regarding treatment with control CM and SASP, 2000 Huh7 cells were seeded in 6-well dishes and treated with fresh aliquots daily for 6 days.

### SA-β-Galactosidase staining and Imaging

Embryos stained for SA-β-Galactosidase (SA-β-gal) activity were initially washed in PBS and then fixed overnight at 4°C using 0.5% glutaraldehyde (Sigma G7651) while agitating gently. The next morning, embryos were washed three times, each 10 minutes in PBS supplemented with 1 mM MgCl2 at a pH of 5.5 and kept at 4°C until further use. The staining process was performed using an X-Gal solution, prepared by dissolving 1 mg/mL X-Gal (Biosynth AG) in PBS (PH 5.5) containing 1 mM MgCl2, along with 5 mM 5 mM K3Fe(CN)6, and 5 mM K4Fe(CN)6 3H2O. The embryos were immersed for 3 to 8 hours until either the AER or the otic vesicles showed positive staining. Embryos were washed again with PBS and then imaged using a Leica MacroFluo Z16 APO macroscope. Regarding Huh7 cells, they were fixed with 0.5% glutaraldehyde for 15 minutes at RT, and washed twice with PBS-MgCl_2_ buffer at pH 6. SA-β-gal staining was done at PH 6 for 5h at 37°C. Imaging was done with an EVOS XL Core (Invitrogen™, RGB images) or an EVOS M5000 (Invitrogen™, Grayscale images).

### Huh7 phenotypic counting

Several phase-contrast images of Huh7 for each condition were taken at random locations (n= 3-9). Colony area expressed in µm² was measured using ImageJ. A “layered colony” was visually determined if a colony was composed of more than one cell layer, either partially or entirely.

### Fluorescence-Activated Cell Sorting (FACS) isolation of AER cells

Forelimb buds were isolated from WT C57Bl/6J or heterozygous p21-mCherry-CreERT2 embryos at E11.5 in cold PBS. Harvested limbs were pooled for enzymatic digestion at 4°C in 2% trypsin (Gibco 15090046) while agitating gently for 5 minutes. Limbs were transferred to PBS with 10% FCS and vortexed vigorously until the ectoderm and mesenchyme were separated. Ectodermal jackets were retained and placed in a new petri dish, where they were subjected to further digestion in 0.05% trypsin at room temperature (RT) with continuous agitation. Jackets were manually dissociated every 5 to 10 minutes using a p200 pipette until full tissue dissociation. The enzymatic reaction was stopped by adding PBS with 10% FCS, and then centrifuged for 5 minutes. The pellet was rinsed with PBS and re-centrifuged before being resuspended in Pre-Sort Buffer (BD Biosciences-563503). The resulting single-cell suspension was filtered through a 50µm nylon mesh filter, kept on ice, and sorted by Flow Cytometry. The gating strategy was established based on mCherry fluorescence levels and negative controls obtained from dissected C57Bl/6J limbs. Sorted cells were collected in Pre-Sort Buffer, centrifuged at 4°C for 5 minutes, rinsed, and re-centrifuged before adding 800 µL TRIzol (Thermofisher scientific 15596026). Cells were subsequently snap-frozen in liquid nitrogen and stored at -80°C until further use.

### RNA extraction

For AER samples, cells previously frozen in TRIzol were thawed on ice and homogenized by vortexing. After a 6-minute incubation at RT, 160µL of chloroform was added and samples were vortexed for 15 seconds before being incubated for an additional 3 minutes at RT. To achieve phase separation, a 12000g centrifugation was performed for 15 minutes at 4°C. The aqueous phase was transferred to a new tube containing 160µL of chloroform. The previous steps were repeated before transferring it to a new empty tube. 1µL of glycogen (20 µg/µL; ThermoFisher, R0551) was added to each sample followed by 400µL of isopropanol. Following a 10-minute incubation at RT samples were centrifugation at 12,000g for another 10 minutes at 4°C. The supernatant was discarded, and the pellets were washed sequentially with 100% ethanol and twice with 75% ethanol. Regarding Huh7 cells, total RNA was extracted from cells using TRIzol, following the manufacturer’s recommendations (Thermofisher scientific, 15596026). Regarding PMK, total RNA was extracted from cells using Direct-zol RNA Miniprep kit (Zymo Research, R2053) and eluted with RNase-free water. For all, after drying the RNA at RT, RNA was resuspended in RNase-free water. RNA concentration was measured using a Nanodrop 2000c spectrophotometer (Thermofisher) and samples were stored at -80°C until further use.

### Sequencing and Meta-analysis

For AER and Huh7 samples, library preparation was done at the GenomEast facility within the IGBMC, following the guidelines provided in the Illumina Stranded mRNA Prep Ligation - Reference Guide (PN 1000000124518). Libraries for RNA sequencing were prepared in accordance with the manufacturer’s guidelines using 40 ng of total RNA, along with the Illumina Stranded mRNA Prep Kit, Ligation Kit, and IDT for Illumina RNA UD Indexes Ligation (all from Illumina, San Diego, USA). Libraries were sequenced on an Illumina NextSeq 2000 sequencer, generating single read 50 base reads.

For AER, PMK and Huh7 samples, preprocessing steps were applied to eliminate adapter sequences, polyA tails, and low-quality reads. Subsequently, reads shorter than 40 bp were eliminated from further analysis. The following preprocessing steps were executed utilizing cutadapt v4.2^42^. Reads were aligned to rRNA sequences from the RefSeq and GenBank databases using bowtie v2.2.8^43^, and aligning sequences were eliminated prior to analysis. The remaining reads were then mapped onto the GRCh38 genome assembly for Homo sapiens or the GRCm39 genome assembly for Mus musculus genome via STAR v2.7.10b^44^. Gene expression quantification was carried out using htseq-count v0.13.5, using annotations from Ensembl v108 and the “union” mode 3^45^. Only unambiguous reads mapping to a specific gene were used for subsequent analysis. Read counts were standardized among samples employing the median-of-ratios method suggested by Anders and Huber, ensuring comparable results across samples^46^. For the remaining senescence datasets, files were downloaded from publicly available Bulk RNA sequencing data deposited either on Gene Expression Omnibus (GEO) or on the European Nucleotide Archive (ENA) as detailed in Extended Data Fig.1a. Regarding datasets extracted from ENA, fastaq files were collected and count files were subsequently generated by the IGBMC sequencing facility using the same pipeline as mentioned above. For datasets published on GEO, available raw count files were used. For each dataset, genes with fewer than 10 raw counts across all samples were excluded from downstream analysis. Normalization was done with DESeq2 version 1.38.1, using Wald test as proposed by Love and Colleagues^47^. Benjamini and Hochberg method was used to adjust P-values for multiple testing.

### Pathway analysis

Gene Set Enrichment Analysis (GSEA) was performed using clusterprofiler (version 4.7.1.002)^48^ with the fast GSEA (fGSEA) algorithm^49^. Over Representation Analysis (ORA) was performed with either clusterprofiler or the gprofiler2 library (version 0.2.3, gprofiler version e113 eg59 p19 f6a03c19)^50^. For each dataset, genes displaying a base mean (average counts across all samples and replicates) of less than 25 bases were excluded prior to analysis. Regarding GSEA, genes were sorted in descending order according to their Wald statistic values, with the top 19,000 genes from the highest to the lowest ranked being used for the analysis. GSEA using fGSEA with a P-value cut-off of 0.05 was performed followed by a Benjamini-Hochberg correction. Minimum and maximum gene set size were limited to between 10 and 2000 genes. Regarding ORA, genes were further filtered, first, using only significant genes with an adjusted P-value below 0.05 and when expressed greater than 1.5 log2 fold, and second before analysis when applicable, utilizing significant genes less than 1.5 log2 fold. For SASP treated PMK, AER and Huh7 samples a ±1.5-fold change threshold was used instead. A cumulative hypergeometric test (one-sided Fisher’s exact test) was used to calculate P-values. P-values were corrected using a BH correction, except for AER and Huh7 samples where the gprofiler2 library was used to perform g:SCS^50^ for multiple testing corrections. The Gene Ontology (GO) term database was used for pathway analysis. Significant terms were determined if corrected P-value < 0.05. Terms pertinent to developmental-related pathways that were significantly enriched or overrepresented, were identified using custom filtering criteria. These included terms containing specific words like “develop”, “regenerate”, “embryonic”, “fate”, “pluripotent”, “layer”, and others. Any irrelevant terms resulting from this were manually excluded.

### Secreted Factors and Transcription factors annotations

Secreted proteins were identified using the Human Protein Atlas database (version 24.0)^21^. Annotated proteins were downloaded from their webpage (URL: https://www.proteinatlas.org/download/proteinatlas.tsv.zip) and filtered for all the proteins with functions related to secretion. The resulting human secretome was converted when necessary to mouse homologs using a homology table provided by the Mouse Genome Database (MGD), (Mouse Genome Informatics (MGI), The Jackson Laboratory, Bar Harbor, Maine. World Wide Web (URL: https://informatics.jax.org/homology.shtml). To minimize ambiguity between both species, we limited the data to one-to-one homologs between human and mouse, where each human gene has exactly one mouse counterpart and the mouse genes also reciprocally only map to the same human gene. This excludes any genes involved in one-to-many or many-to-many relationships. Except for some cases where a gene has a one-to-many homolog relationship but shares a same gene symbol in both human and mouse annotations, in that case the identically named homolog was also retained in the final gene list. Finally, to ensure consistency and facilitate gene symbol usage across species whenever both human and mouse data were presented at the same time, genes from both species were shown using only the HGNC human gene symbols, with the corresponding mouse homologs relabeled accordingly. AnimalTFDB version 4.0491^51^ was used to annotate transcription factors for both mouse and human data. Data manipulation was mostly done using R (version 4.2.2 to 4.5.1). Most graphs were generated using ggplot2 (up to version 3.5.2). Transcription factor bar graphs and Huh7 cell counting graphs were created with Prism (GraphPad, version 10.6.0).

### Data availability

The omics datasets generated during this study (AER, PMK, Huh7) will be deposited in the GEO database and made publicly available before publishing. Remaining data is available in the main text, in the supplementary materials or publicly available as indicated in the methods. Custom R scripts, along with any intermediary files used for data generation, manipulation, and analysis, are freely accessible upon reasonable request.

## ACKNOWLEDGEMENTS

This work was supported by grants from La Fondation pour la Recherche Medicale (FRM) Amorcage pour les jeunes equipes (AJE20160635985); Fondation ARC pour la Recherche sur le Cancer (PJA20181208104); Agence Nationale de la Recherche (ANR) ANR-19-CE13-0023 and ANR-22-CE14-0062; Ligue Contre le Cancer - Equipe Labellisée 2024, and Institut National du Cancer (INCA-PLBIO23-004-2023-134) (all to W.M. K.). D.S.G was supported by the National Research Fund, Luxembourg (FNR) (AFR PhD, Application ID:14584624) and the University of Strasbourg. A.K was supported by a fellowship from Eur IMCBio – University of Strasbourg and the Fondation ARC pour la Recherche sur le Cancer, France fourth-year PhD fellowship. L.D is funded by an École Doctorale fellowship, University of Strasbourg, France. The work was also supported by an institutional grant of the Interdisciplinary Thematic Institute IMCBio+, as part of the ITI 2021-2028 program of the University of Strasbourg, CNRS and Inserm, was supported by IdEx Unistra (ANR-10-IDEX-0002), and by SFRI-STRAT’US project (ANR-20-SFRI-0012) and EUR IMCBio (ANR-17-EURE-0023) under the framework of the France 2030 Program. The French National Research Agency (ANR) supported the generation, breeding, and phenotyping of mutant mice reported here through the Programme d’Investissement d’Avenir under contract INBS PHENOMIN (ANR-10-INBS-07) grant under the frame programme Investissement d’Avenir ANR-10-IDEX-0002-02. Sequencing was performed by the GenomEast platform, a member of the “France Genomique” consortium (ANR-10-INBS-0009). The funders had no role in study design, data collection and analysis, decision to publish, or preparation of the manuscript.

## SUPPLEMENTARY TABLES

**Supplementary Table 1. Upregulated secreted factors in the AER.** Secreted Factors upregulated more than 1.5-fold in the AER compared to Ectoderm cells.

**Supplementary Table 2. Secreted factors upregulated in senescence.** Secreted Factors significantly upregulated (>25 normalized counts, adjusted P <0.05; >1.5-fold) across different senescence datasets.

**Supplementary Table 3. Muscle transcriptome in senescence.** Differentially expressed genes (>25 normalized counts) in muscle tissue comparing senescent to non-senescent cells under various conditions.

**Supplementary Table 4. Upregulated genes coding for secreted proteins in senescent satellite cells.** Significantly upregulated secreted factors (adjusted P < 0.05; >1.5-fold change) in senescent satellite cells post-injury.

**Supplementary Table 5. SASP-induced gene expression changes in primary mouse keratinocytes.** Differentially expressed genes in primary mouse keratinocytes following treatment with SASP for 2 or 6 days compared to control CM.

**Supplementary Table 6. Differentially expressed transcription factors in primary mouse keratinocytes.** Differentially expressed transcription factors in SASP treated primary mouse keratinocytes for 2 or 6 days compared to control CM.

**Supplementary Table 7. Gene expression in doxorubicin-induced senescence in Huh-7 cells.** Differentially expressed genes in doxorubicin induced senescence in Huh-7 cells, compared to non-treated proliferating Huh-7 cells.

**Supplementary Table 8. Secreted factors in doxorubicin-induced senescence in Huh**-**7 cells.** Significantly upregulated secreted factors (adjusted-P <0.05; >1.5-fold) in doxorubicin-treated Huh-7 cells.

**Supplementary Table 9. GO terms related to development, from analysis of secreted factors in doxorubicin induced senescence.** Overrepresented GO terms related to development, from analysis of upregulated secreted factors in doxorubicin induced senescence in Huh-7 cells.

**Supplementary Table 10. Gene expression in Huh-7 cells after exposure to the SASP.** Differentially expressed genes in Huh-7 cells following treatment with SASP for 6 days compared to Huh-7 cells treated with control CM.

**Supplementary Table 11. Upregulated TFs in SASP-treated Huh-7 cells.** Significantly upregulated (adjusted P <0.05; >0 fold) transcription factors in SASP treated Huh-7 cells.

---

**Extended Data Fig. 1.**
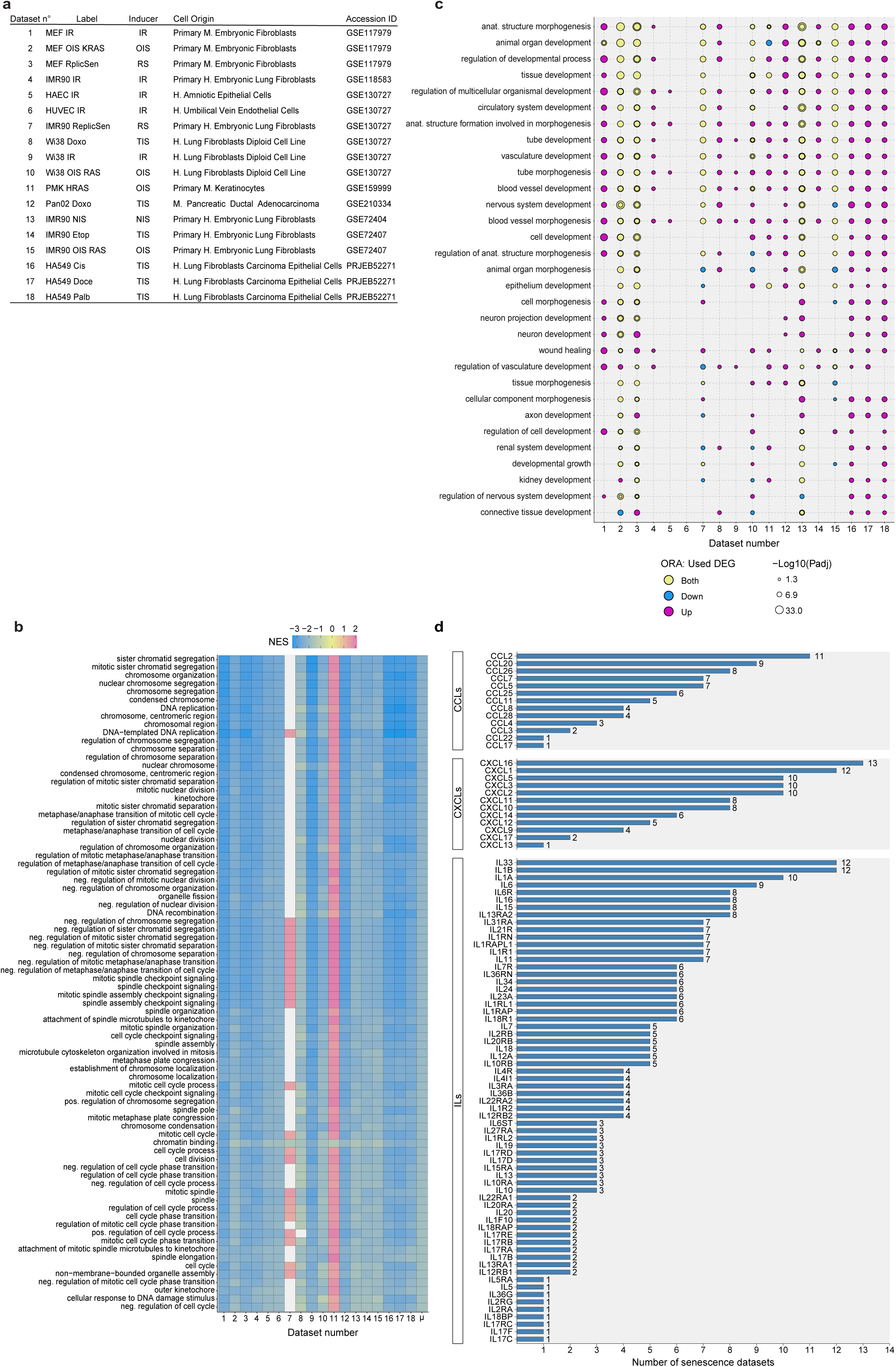
Gene-expression analysis confirms an enrichment of senescence related biological processes, the SASP, and developmental signatures in most senescent datasets. **(a)** The 18 different senescence models used for comparing bulk RNA sequencing datasets. IR: Irradiation, OIS: Oncogene-induced senescence, RepliSen: Replicative senescence, Doxo: Doxorubicin, NIS: Notch induced senescence, Etop: Etoposide, Cis: Cisplatin, Doce: Docetaxel, Palb: Palbociclib, KRAS: Kirsten rat sarcoma virus, HRAS: Harvey Rat sarcoma virus protein. (**b**) GSEA was performed on every dataset separately, and the top 80 terms that were negatively enriched on average and present in at least 17 datasets were selected and plotted. The X-axis shows the different datasets, with tiles representing their individual normalized enrichment score (NES) per term. ‘µ’ represents the average NES across every dataset and was used to sort the terms in an ascending order. (**c**) Overrepresentation analysis (ORA) showing significantly overrepresented GO terms related to development, present in 7 or more datasets. Analysis performed using genes that were significantly up- or down-regulated (adjusted P < 0.05; >/<1.5 log_2_ fold, exceeding 25 normalized counts on average). (**d**) Bar graph showing a subset of Interleukins (ILs), CC Motif Chemokine Ligands (CCLs) and CXC Motif Chemokine Ligands (CXCLs), some of which are known to be differentially expressed in senescence. X-axis represents the number of senescent datasets in which the selected genes are significantly (adjusted P < 0.05) upregulated (>1.5-fold) in senescent cells.

**Extended Data Fig. 2.**
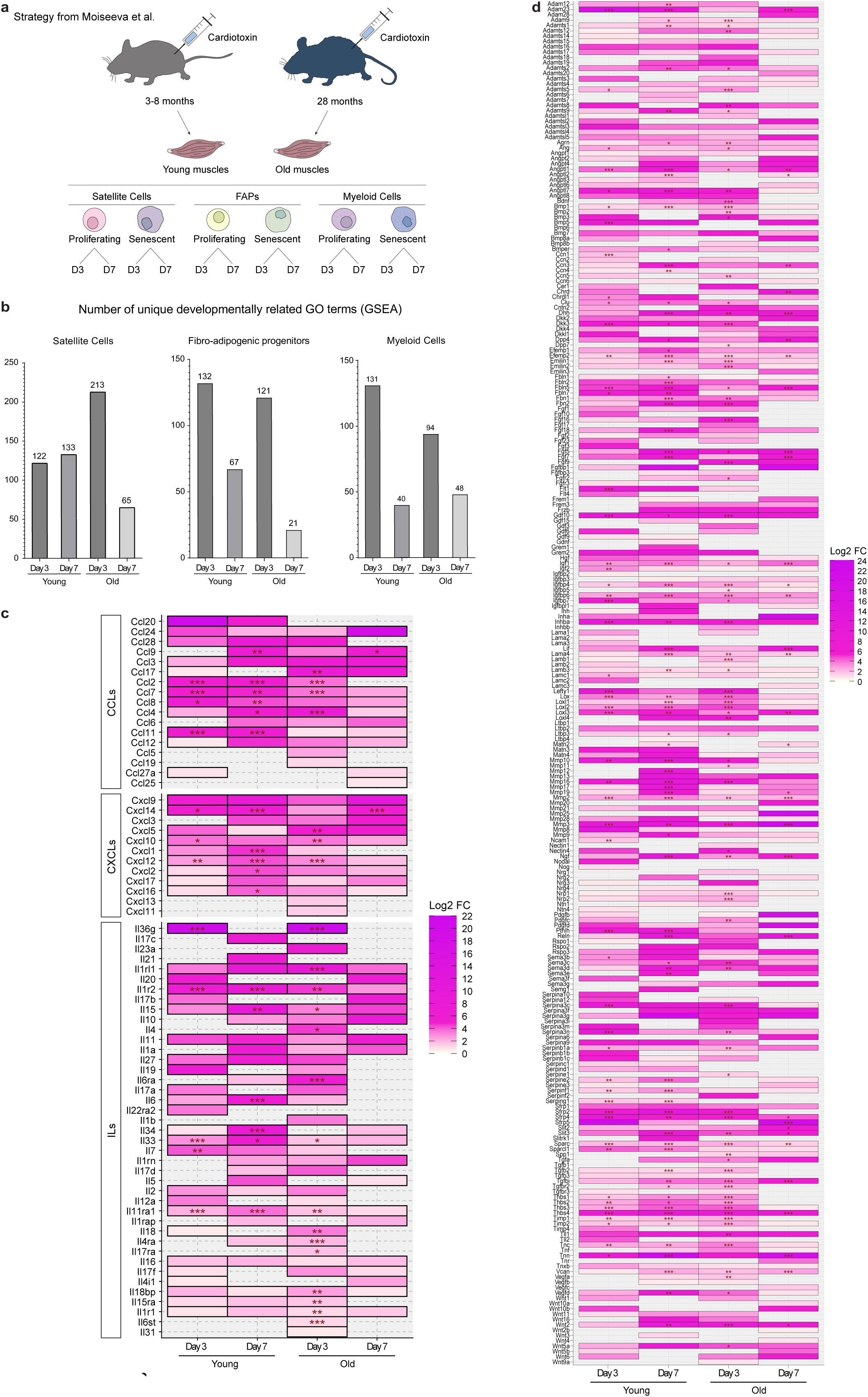
RNA-seq analysis on senescent cells isolated from muscle. **(a)** A schematic illustrating the strategy from Moiseeva et al. to select muscle cells for RNA sequencing. The authors collected muscle from 3- to 8-months old (= young) mice and 28 months old (= old) mice, 3- and 7-days post-injury via cardiotoxin injection. Muscle cells were sorted into different cell types, differentiating between SA-ß-Gal negative and positive cells prior to sequencing. (**b**) Total number of significantly enriched Gene Ontology (GO) terms identified by GSEA in the different senescent muscle cells, when compared to their individual non-senescent counterparts, corresponding to developmentally related biological processes. (**c**) Heatmap representing pattern of expression in satellite cells, of genes coding for secreted proteins belonging to the Interleukin (ILs), CC Motif Chemokine Ligands (CCLs) and CXC Motif Chemokine Ligands (CXCLs) families. (**d**) Similar Heatmap as in (c) but with selected secreted factors associated with developmental processes. Significant differently expressed genes are marked as follows: * (adjusted P < 0.05), ** (adjusted P < 0.01), and *** (adjusted P < 0.001).

**Extended Data Fig. 3.**
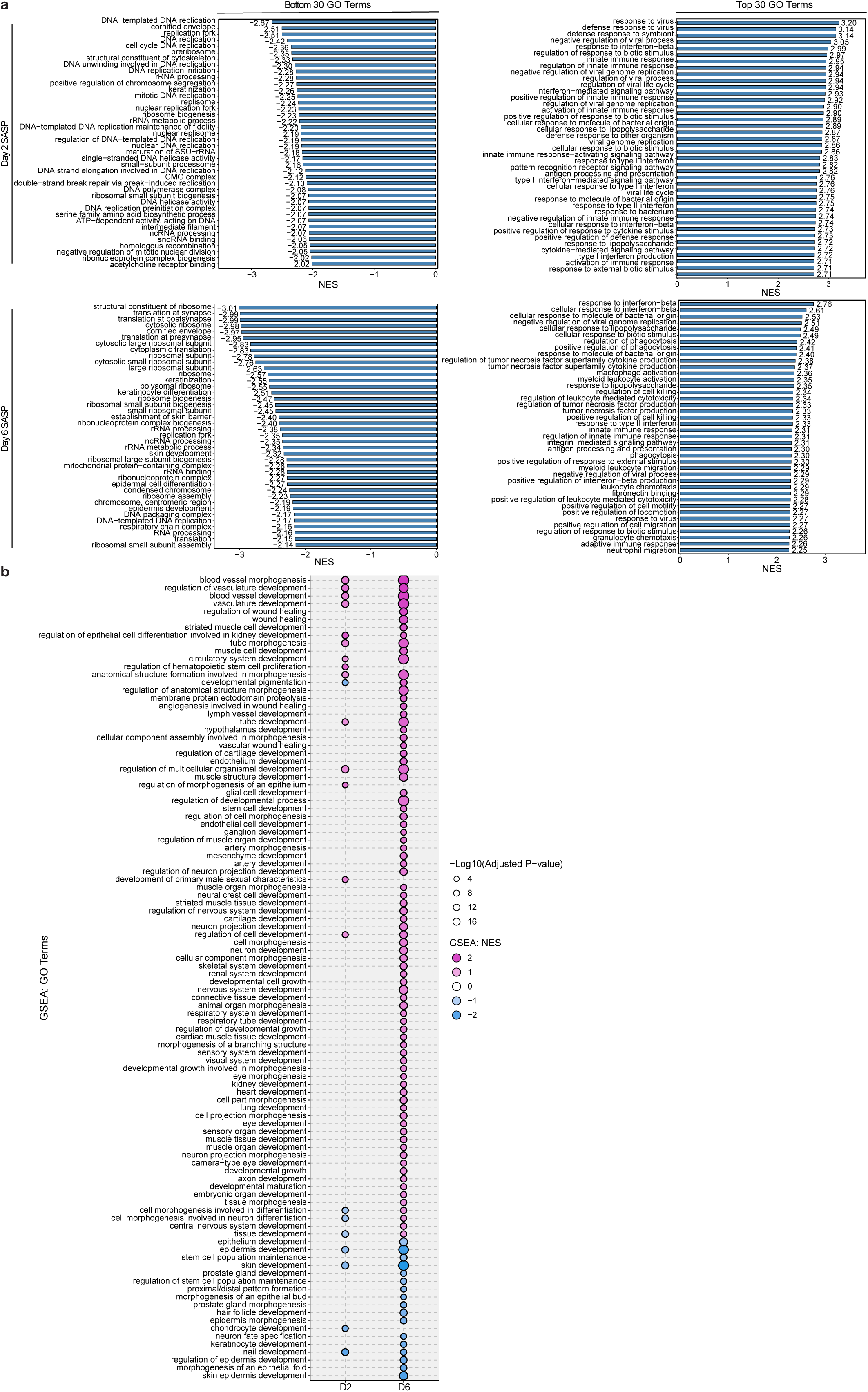
RNA-seq analysis of primary mouse keratinocytes (PMK) exposed to the SASP from OIS in PMK. **(a)** Gene Ontology (GO) terms significantly (P < 0.05) enriched by gene set enrichment analysis (GSEA) in Primary Mouse Keratinocytes (PMK) treated with SASP for either 2x24h (Day 2) or 6x 24h (Day 6) compared to control conditioned media (CM). Left bar graphs display the top 30 most negatively enriched GO terms, as indicated by a negative Normalized Enrichment Scores (NES). Graphs on the right illustrate the top 30 most positively enriched GO terms. (**b**) GO terms related to development, that were significantly enriched by GSEA in PMK following treatment with SASP for 2 or 6 days compared to control CM.

## Notes

### Competing Interest Statement

The authors have declared no competing interest.

### Summary of Updates

-cross-referencing another BioRxiv preprint -minor text changes

